# The more synthetic polymer types pollute the soil, the stronger the growth suppression of invasive alien and native plants

**DOI:** 10.1101/2022.12.08.519663

**Authors:** Yanmei Fu, Mark van Kleunen, Kai Ma, Yanjie Liu

## Abstract

Although most studies on the ecological effects of microplastic pollution focus on a single type of synthetic polymer and a single species, most organisms will be exposed to multiple polymer types simultaneously and the effects may vary among species. To test the effects of polymer diversity on plants, we grew single plants of eight invasive and eight native species in pots with substrate polluted by 0, 1, 3 and 6 types of micro-sized synthetic polymers. We found that the growth suppression by microplastic pollution became stronger with the number of polymer types the plants were exposed to. This tended to be particularly the case for invasive species, whose biomass advantage over natives diminished with the number of polymer types. Our study thus shows that the negative effects of microplastic pollution on plant growth increase with the number of polymer types, and that these effects differ between invasive and native species.

## Introduction

Microplastic pollution is a prominent characteristic of the Anthropocene (Barnes *et al*. 2009; de Souza Machado *et al*. 2018), but research on it has long been ignored until the term ‘microplastic’was prosed in 2004 (Thompson *et al*. 2004). As microplastic pollution could potentially affect many biological processes from cellular to ecosystem levels (Baho, Bundschuh & Futter 2021), it has rapidly become a hot topic in ecological research. However, although terrestrial ecosystems are a major sink of microplastics (Jambeck *et al*. 2015; Xu *et al*. 2020; Baho, Bundschuh & Futter 2021), aquatic systems have received much more research attention than terrestrial ones.

Given the prevalence of microplastics in many soils (Rillig & Lehmann 2020), exposure of terrestrial plants to microplastics is inevitable. Microplastic pollution is hypothesized to affect plant performance via various mechanisms, such as direct toxicity and alterations of the soil environment (Rillig *et al*. 2019a). For example, Bosker *et al*. (2019) found that microplastic particles of the synthetic polymer polystyrene delayed germination and root growth of *Lepidium sativum* due to physical blockage of the pores in the seed capsule. An experiment by de Souza Machado *et al*. (2019) revealed that microplastic particles made of polyester decreased soil bulk density, and increased water saturation and microbial activity, which resulted in increased biomass production of *Allium fistulosum*. These and other recent studies on the potential effects of microplastics on terrestrial plants (Boots, Russell & Green 2019; Sun *et al*. 2020; Lozano *et al*. 2021a; Yang *et al*. 2021; Luo *et al*. 2022) have mainly focused on the ecological effects of single specific synthetic polymers on plants. However, plastics polluting the soil come from various synthetic polymers used in everyday life (Geyer, Jambeck & Law 2017; Wei *et al*. 2021). Yet, little is known about how plants respond to microplastic pollution caused by more than one type of polymer simultaneously.

The studies on microplastic pollution and plants indicate that the various polymers differ in their individual effects on plant growth (de Souza Machado *et al*. 2018; van Kleunen *et al*. 2020; Lozano *et al*. 2021b; Meng *et al*. 2021). There are different ways in which the combined effects of multiple polymer types could deviate from the additive effects of the individual polymers. On the one hand, the negative effects of microplastics may increase with diversity of polymers because, with an increasing number of polymer types, there is a higher probability that one of them has a strong negative effect (e.g. toxic effect) on plants (i.e. similar to a sampling effect in biodiversity ecosystem experiments; Loreau & Hector 2001) or that some of the polymers act synergistically rather than additively (Figure 1a). On the other hand, if the total amount of microplastics remains the same, a higher diversity of polymers may dilute the individual polymer types to levels that do not affect plant growth (Figure 1b). Which one of these two scenarios is more likely requires empirical tests with many different polymer types and plant species.

**Figure 1.**
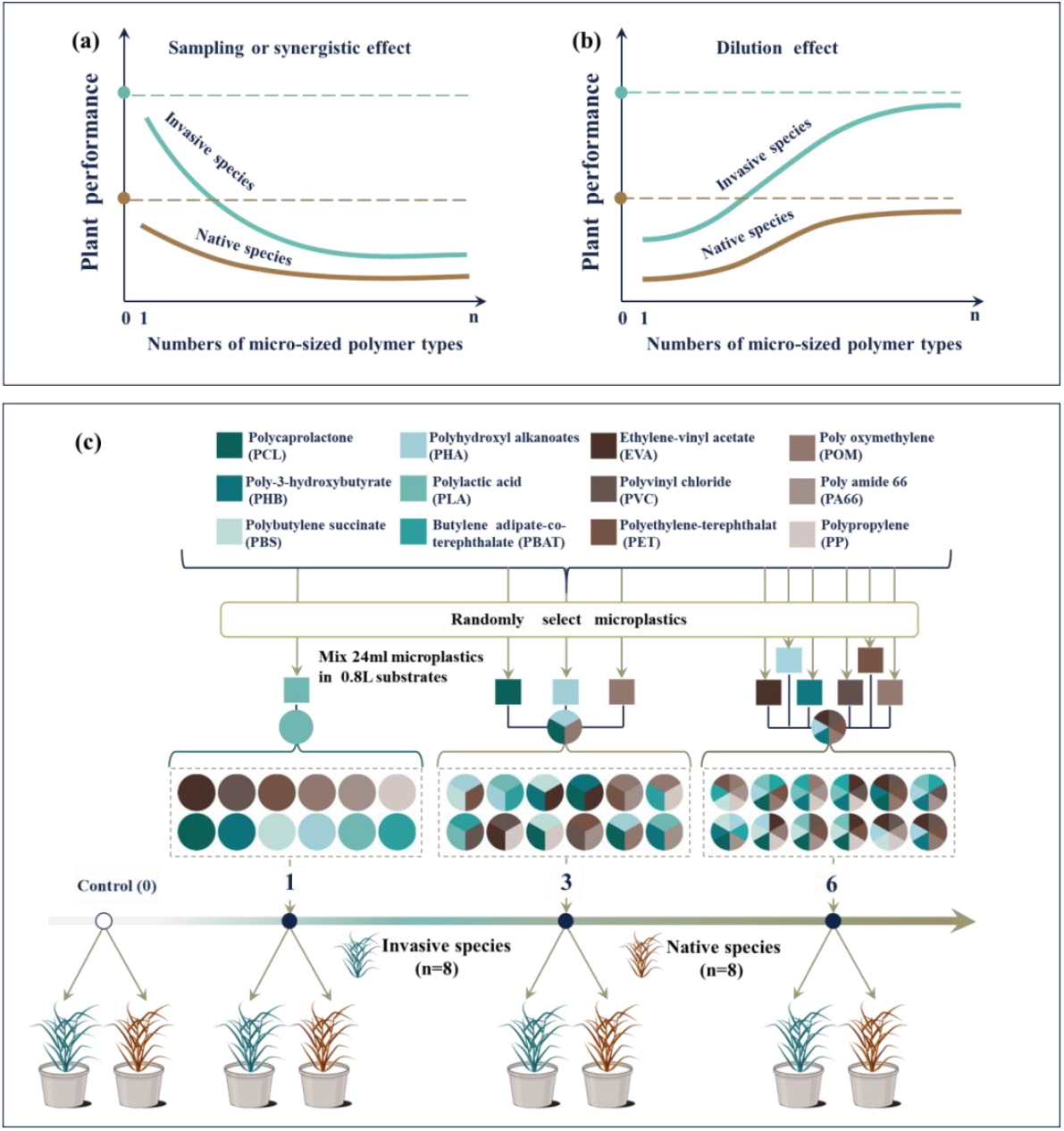
Diagrams illustrating the predictions of how increased numbers of micro-sized polymer types could affect growth of invasive and native species, as well as the experimental design testing the predictions. (a) Graphical representation of the hypothesis that increasing numbers of polymer types will result in stronger negative effects on plants due to sampling or synergistic effects, and how this may differ between invasive alien and native plants; (b) Graphical representation of the hypothesis that increasing numbers of polymer types will result in weaker negative effect on plants due to dilution effects, and how this may differ between invasive alien and native plants. (c) A schematic illustration of the experimental design.

The invasion by alien species is another key characteristic of the Anthropocene (Lewis & Maslin 2015). Alien plant species have invaded almost all regions of the world (van Kleunen *et al*. 2015), and their numbers are still increasing (Seebens *et al*. 2018). Alien plants are predominantly found in anthropogenic habitats (Chytrý *et al*. 2008), which are also likely the habitats most strongly exposed to plastic pollution. Whether microplastics affect alien and native plant species differently, however, has rarely been tested (but see Lozano & Rillig 2020). Richards et al. (2006) proposed that phenotypic plasticity may allow successful invaders to maintain fitness in stressful environments, to take advantage of less stressful environments or a combination of both abilities. Which one of those scenarios applies with regard to the responses of alien and native plants to the number of polymer types remains to be tested.

To test the effects of microplastic pollution and increasing numbers of micro-sized polymer types in the soil on alien and native plants, we conducted a multi-species greenhouse experiment. We grew single plants of eight invasive alien and eight native species in pots with substrate polluted by 0, 1, 3 and 6 types of micro-sized polymers. By comparing total biomass production, maximum height, and root morphology and allocation of the two groups of plant species in the different microplastic-pollution treatments, we addressed the following specific questions: (i) Does microplastic pollution have a negative effect on plant growth, and does this differ between invasive alien and native plants? (ii) Does the diversity of micro-sized polymer types result in stronger or weaker effects on plant growth, and does this differ between invasive alien and native plants?

## Material and Methods

### Study species

To increase the generality of our findings, we selected a total of 16 herbaceous species belonging to five families (Table S1). All 16 species are common in China, and we classified each species as invasive alien or native to China based on the *Alien Invasive Flora of China* (Ma 2020) and the online database Flora of China (www.efloras.org). Seeds of each species were collected and mixed from at least four natural grassland populations (at least 1 km apart).

### Microplastic pool

To simulate microplastic pollution by different numbers of polymer types, we used a pool of 12 types of commonly used synthetic polymers: polybutylene succinate (PBS; Mulvane, Kansas, USA), polycaprolactone (PCL; Malmö, Sweden), polyhydroxyl alkanoates (PHA; Shenzhen, China), poly-3-hydroxybutyrate (PHB; Rialto, California, USA), polylactic acid (PLA; Willich, Germany), polybutylene adipate-co-terephthalate (PBAT; Kansas, USA), ethylene-vinyl acetate (EVA; Seoul, South Korea), polyamide 66 (PA66; Wilmington, Delaware, USA), polyethylene-terephthalat (PET; Barcelona, Spain), poly oxymethylene (POM; Wilmington, Delaware, USA), polyvinyl chloride (PVC; Leominster, Massachusetts, USA) and polypropylene (PP; Lisboa, Portugal). All polymers were acquired in powder form. As the 12 ready-made microplastics (i.e. powders) were acquired from different commercial suppliers, the sizes of the polymer particles differed as the producers either had passed them through sieves with a mesh width of 150 µm (PCL, PHA, PHB, PLA, PVC, POM, PP, PA66) or 180 µm (PBS, PBAT, EVA, PET). We purchased these microplastics from Suzhou XinSuYu Company in Suzhou, China.

### Experimental set-up

The experiment was performed in a greenhouse of the Northeast Institute of Geography and Agroecology, Chinese Academy of Sciences (43°59’49‖ N, 125°24’03‖E). From 27 August to 17 September 2021 (Table S1), we sowed the seeds of each species into separate trays (20cm × 20cm × 4.5cm) filled with commercial potting soil (Pindstrup Plus, Pindstrup Mosebrug A/S, Ryomgård, Denmark). Since the 16 species were known to have different germination times (Yanmei Fu, personal observation), we sowed them on different dates to ensure that all were at a similar developmental stage at transplanting. On 18 September 2021, we selected 48 similar-sized seedlings of each species, and transplanted them individually into the centre of 1L circular plastic pots (i.e. one individual per pot) filled with 0.8 L substrate. For the control pots without microplastic pollution, the substrate was a 1:1 (v:v) mixture of washed sand and fine vermiculite. For the pots with microplatic pollution, the substrate was a 97:97:6 (v:v:v) mixture of washed sand, fine vermiculite and polymer powder. To ensure that the substrate contained enough nutrients for plant growth, we also mixed 4 g of a slow-release fertilizer (Osmocote Exact Standard, Everris International B.V., Geldermalsen, Netherlands) in the substrate of each pot.

The microplastic-pollution treatment included four levels, i.e. no pollution and pollution with one, three or six types of polymer. Each level of the treatment was replicated 12 times per plant species. For the pollution level with one type of polymer, we used each of the 12 polymers once (i.e. PBS, PCL, PHA, PHB, PLA, PBAT, EVA, PA66, PET, POM, PVC and PP; Figure 1c). For the pollution level with three types of polymers, we made 12 different combinations of polymers with the restriction that each polymer type was used in three different combinations (Figure 1c; Table S2). For the pollution level with six types of polymers, we also made 12 different combinations of polymers but with the restriction that each polymer type was used in six different combinations (Figure 1c; Table S2). For each pot of the three polymer-diversity levels, we mixed 0.024 L powdered polymer into the substrate (i.e. 3% volume of total substrate). In other words, we mixed 0.024 L of each selected polymer into pots with one type polymer, 0.008 L of each selected polymer into pots with three types of polymer, and 0.004 L of each selected polymer into pots with six types of polymer (Figure 1c). The concentration of polymers that we mixed into the substrate is within the range of microplastic pollution reported for soils in terrestrial ecosystems (Fuller & Gautam 2016; Rillig 2018). In total, the experiment included 768 pots: 2 status (alien *vs* native) × 8 species per status × 4 levels of the microplastic-pollution treatment (0, 1, 3, 6 types of polymer) × 12 replicates. All pots were randomly distributed on four benches of a greenhouse with a temperature between 22°C and 28°C and a 13 h light/11 h dark cycle. We re-randomized positions of all pots after four weeks (i.e. 16 October 2021). To ensure that there was no water limitation during the experiment, we regularly watered the plants.

### Measurements

On 20 September 2021, three days after transplanting, we measured the initial heights of all plants. On 24 November 2021, before plant harvest, we measured the final plant height of each plant. We then harvested for each plant the aboveground and belowground biomass separately. The aboveground biomass of each plant was directly dried at 65°C for 72 h, and weighed. After washing the roots of each plant free from substrate, we cut them into 2cm fragments. We then randomly picked a representative subsample from each individual root system, and determined the root length and the mean root diameter using a flatbed scanner specifically modified for root scanning (Epson Expression 12000 XL; Seiko Epson Corporation [SEKEY), Japan) and WinRhizo software (version 2019a; Regent Instruments Inc., Québec City, QC, Canada). All scanned root subsamples, as well as the remaining root mass of each plant were also dried for at least 72 hours at 65°C, and weighed. Because ten plants died during the experiment, we only harvested 758 instead of 768 plants.

Based on final biomass and root morphology measurements, we calculated total biomass (aboveground + belowground biomass), root mass fraction (belowground biomass / total biomass), total root length (root length of subsample × total belowground biomass / subsample biomass) and specific root length (root length of subsample / subsample biomass).

### Statistical analysis

All statistical analyses were conducted using R 4.1.0 (R Core Team, 2020). To test whether and how the presence of microplastic pollution affects growth performance, root allocation and morphology of alien and native plants, we fitted linear mixed-effect models using the function lme in the ‘nlme’ package (Pinheiro et al. 2020). In the models, total biomass, final plant height, root mass fraction, root diameter and specific root length were response variables. We included the microplastic pollution (presence [i.e. averaged across 1, 3 and 6 polymer types] *vs* absence [i.e. 0 polymer type])), invasion status (invasive *vs* native) and their interaction as fixed effects in each of above models. To test whether and how the diversity of micro-sized polymer types affects growth performance, root allocation and morphology of alien and native plants, we also fitted linear mixed-effect models for the subset of plants grown in the presence of microplastics, in which total biomass, final plant height, root mass fraction, root diameter and specific root length were response variables. In these models, we included the diversity of micro-sized polymer types (1, 3 *vs* 6), invasion status (invasive *vs* native) and their interaction as fixed effects.

For all above models, total biomass, root diameter and specific root length were natural-log-transformed, final plant height was square-root-transformed, and root mass fraction was logit-transformed to meet the assumptions of normality. To control for potential effects of initial plant size on final plant performance, we also included initial plant height as scaled natural-log-transformed covariate in all models. To account for the non-independence of individuals of the same species and the phylogenetic non-independence of the species, we included species nested within family as random effects in all models. In addition, to account for the non-independence of replicates of microplastic-pollution levels, we also included identity of each microplastic type or mixture as random effect in all models. Due to violation of the homoscedasticity assumption (Zuur *et al*. 2009) of models analyzing total biomass, final plant height and root mass fraction, we allowed the species and microplastic type or composition (Table S2) to have different variances by using the varComb and varIdent functions in the R package ‘nlme’ (Pinheiro et al. 2020). For models analyzing specific root length and root diameter, we also included the variance structure to model different variances per species using the varIdent function. We assessed the significance of fixed effects and their interaction in each model with likelihood-ratio tests (type II) using the Anova function in the R package car (Fox & Weisberg 2019).

## Results

Averaged across all microplastic treatments, invasive alien species produced more total biomass (+90.0%) than native species (Table 1 & 2; Figure 2). Invasive alien species also showed greater specific root length (+33.6%) than native species (Table 1 & 2; Figure 3). Averaged over all 16 species, the presence of microplastic pollution (i.e. averaged across 1, 3 and 6 polymer types) did not affect any traits we measured (Table 1, Figure 2 & 3). However, the presence of microplastic pollution decreased the total biomass of invasive alien species more than those of native species (Table 1; Figure 2). In addition, increased polymer diversity (i.e. from 1 to 6 polymer types) significantly inhibited total biomass (slope, -0.190; Table 2; Figure 2) and specific root length (slope, -0.055; Table 2; Figure 3) of plants, but enhanced root mass fraction (slope, 0.029; Table 2; Figure 3) and root diameter (slope, 0.022; Table 2; Figure 3). Interestingly, both the suppression on specific root length (slope, -0.071 *vs* -0.036; Table 2; Figure 3) and promotion on root diameter (slope, 0.029 *vs* 0.014; Table 2; Figure 3) of plants were stronger for invasive alien species than for native species. Similarly, the suppression of total biomass of plants tended to be stronger for invasive alien species (slope, -0.217) than for native species (slope, -0.172; marginally significant S × PD in Table 2; Figure 3).

**Table 1.** Effects of microplastic pollution (presence [i.e. averaged across 1, 3 and 6 polymer types] *vs* absence [i.e. 0 polymer type]), invasion status (invasive *vs* native) and their interactions on total biomass, final plant height, root mass fraction, specific root length and root diameter of plant species. Significant effects (*p* < 0.5) are highlighted in bold, and marginally significant effects (0.05 < *p* < 0.1) are underlined.

| Terms | Total biomass<br>(natural-log-transformed) |  |  | Plant height<br>(square-root-transformed) |  |  | Root mass fraction<br>(logit-transformed) |  |  | Specific root length<br>(natural-log-transformed) |  |  | Root diameter<br>(natural-log-transformed) |  |  |
| --- | --- | --- | --- | --- | --- | --- | --- | --- | --- | --- | --- | --- | --- | --- | --- |
| | *df* | $\chi^2$ | *p* | *df* | $\chi^2$ | *p* | *df* | $\chi^2$ | *p* | *df* | $\chi^2$ | *p* | *df* | $\chi^2$ | *p* |
| **Fixed terms** |  |  |  |  |  |  |  |  |  |  |  |  |  |  |  |
| Initial height | 1 | 33.737 | **<0.001** | 1 | 8.269 | **0.004** | 1 | 13.488 | **<0.001** | 1 | 0.733 | 0.392 | 1 | 0.008 | 0.927 |
| Status (S) | 1 | 10.480 | **0.001** | 1 | 5.569 | **0.018** | 1 | 1.145 | 0.285 | 1 | 4.705 | **0.030** | 1 | 2.534 | 0.111 |
| Microplastic pollution (MP) | 1 | 0.941 | 0.332 | 1 | 0.879 | 0.348 | 1 | 0.895 | 0.344 | 1 | 0.699 | 0.403 | 1 | 0.907 | 0.341 |
| S × MP | 1 | 13.434 | **<0.001** | 1 | 3.392 | <u>0.065</u> | 1 | 2.364 | 0.124 | 1 | 1.774 | 0.183 | 1 | 0.060 | 0.806 |
| **Random terms** |  |  |  |  |  |  |  |  |  |  |  |  |  |  |  |
|  | SD |  |  | SD |  |  | SD |  |  | SD |  |  | SD |  |  |
| Species | 0.581* |  |  | 1.032* |  |  | 0.599* |  |  | 0.589* |  |  | 0.171* |  |  |
| Family | 1.015 |  |  | 0.937 |  |  | 0.0003 |  |  | 0.397 |  |  | 0.204 |  |  |
| Microplastic type or composition | 1.193* |  |  | 0.978* |  |  | 0.150* |  |  | 0.254 |  |  | 0.109 |  |  |
| | $R^2_m$ | $R^2_c$ | | $R^2_m$ | $R^2_c$ | | $R^2_m$ | $R^2_c$ | | $R^2_m$ | $R^2_c$ | | $R^2_m$ | $R^2_c$ | |
| **$R^2$ of the model** | 0.167 | 0.977 |  | 0.154 | 0.930 |  | 0.071 | 0.888 |  | 0.144 | 0.849 |  | 0.063 | 0.828 |  |

\* Standard deviations for individual species effect for the saturated model are found in TableS3. $R^2_m$ : marginal $R^2$ , $R^2_c$ : conditional $R^2$ .

**Figure 2.**
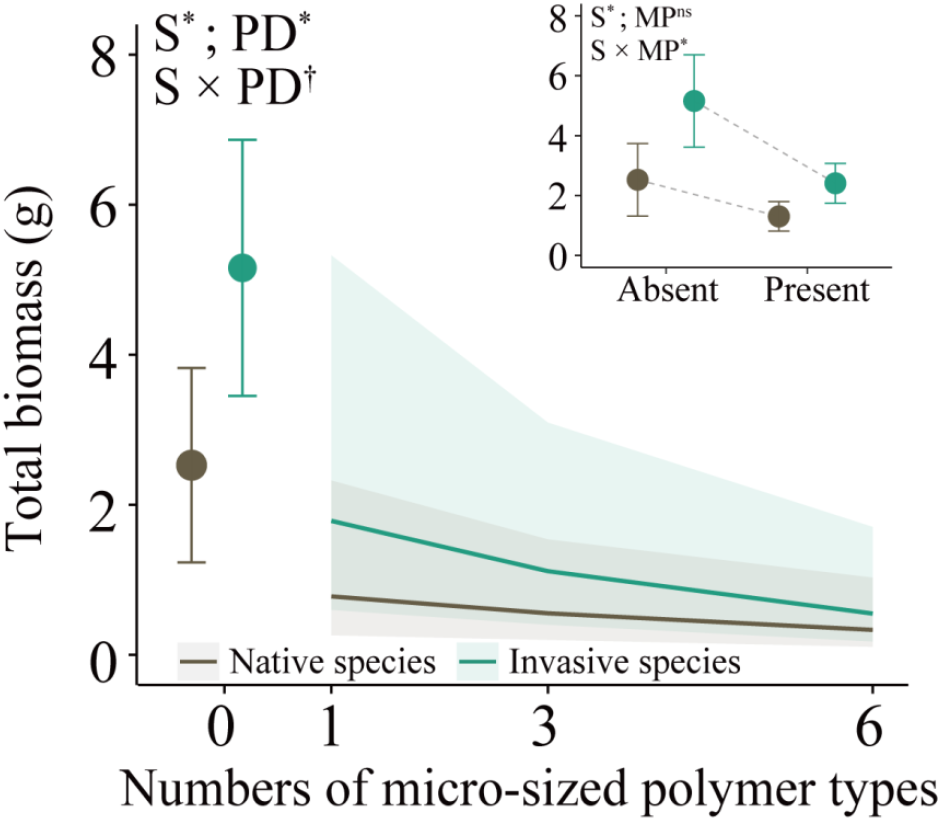
Total biomass of invasive and native species under the different numbers of micro-sized polymer types (0, 1, 3, 6) mixed into the substrate. In the main graph, points with error bars represent mean values ± SE in the absence of microplastic, and solid lines are the fitted relationships for the effect of polymer diversity, and the colored polygons delimit the 95% confidence intervals. Inset at the right upper corner of the graph shows the effects of the absence *vs* presence of microplastic (averaged over 1, 3 and 6 polymer types). Significant parameters (i.e. ‘S’ [invasion status], ‘PD’ [polymer diversity], ‘MP’ [microplastic pollution], ‘S × PD’ and ‘S × MP’) are indicted with asterisks (*), marginally significant ones are indicated with daggers (†), and non-significant ones are indicated with ‘ns’.

**Table 2.** Effects of polymer diversity (1, 3, 6; continuous variable), invasion status (invasive *vs* native) and their interactions on total biomass, final plant height, root mass fraction, specific root length and root diameter of plant species. Significant effects (*p* < 0.5) are highlighted in bold, and marginally significant effects (0.05 < *p* < 0.1) are underlined.

| Terms | Total biomass<br>(natural-log-transformed) |  |  | Plant height<br>(square-root-transformed) |  |  | Root mass fraction<br>(logit-transformed) |  |  | Specific root length<br>(natural-log-transformed) |  |  | Root diameter<br>(natural-log-transformed) |  |  |
| --- | --- | --- | --- | --- | --- | --- | --- | --- | --- | --- | --- | --- | --- | --- | --- |
| | *df* | $\chi^2$ | *p* | *df* | $\chi^2$ | *p* | *df* | $\chi^2$ | *p* | *df* | $\chi^2$ | *p* | *df* | $\chi^2$ | *p* |
| **Fixed terms** |  |  |  |  |  |  |  |  |  |  |  |  |  |  |  |
| Initial height | 1 | 24.885 | **<0.001** | 1 | 5.554 | **0.018** | 1 | 8.928 | **0.003** | 1 | 0.273 | 0.601 | 1 | 0.739 | 0.390 |
| Status (S) | 1 | 7.167 | **0.007** | 1 | 6.144 | **0.013** | 1 | 1.090 | 0.296 | 1 | 4.192 | **0.041** | 1 | 2.396 | 0.122 |
| Polymer diversity (PD) | 1 | 4.413 | **0.036** | 1 | 4.411 | **0.036** | 1 | 5.072 | **0.024** | 1 | 7.130 | **0.008** | 1 | 6.450 | **0.011** |
| S × PD | 1 | 2.982 | <u>0.084</u> | 1 | 3.136 | <u>0.077</u> | 1 | 1.935 | 0.164 | 1 | 3.890 | **0.049** | 1 | 6.841 | **0.009** |
| **Random terms** |  |  |  |  |  |  |  |  |  |  |  |  |  |  |  |
|  | SD |  |  | SD |  |  | SD |  |  | SD |  |  | SD |  |  |
| Species | 0.616* |  |  | 0.936* |  |  | 0.574* |  |  | 0.602* |  |  | 0.172* |  |  |
| Family | 0.983 |  |  | 1.016 |  |  | 0.003 |  |  | 0.406 |  |  | 0.216 |  |  |
| Microplastic type or composition | 1.102* |  |  | 0.907* |  |  | 0.113* |  |  | 0.229 |  |  | 0.101 |  |  |
| | $R^2_m$ | $R^2_c$ | | $R^2_m$ | $R^2_c$ | | $R^2_m$ | $R^2_c$ | | $R^2_m$ | $R^2_c$ | | $R^2_m$ | $R^2_c$ | |
| **$R^2$ of the model** | 0.114 | 0.772 |  | 0.080 | 0.510 |  | 0.012 | 0.141 |  | 0.131 | 0.821 |  | 0.057 | 0.798 |  |

\* Standard deviations for individual species effect for the saturated model are found in Table S4. $R^2_m$ : marginal $R^2$ , $R^2_c$ : conditional $R^2$ .

**Figure 3.**
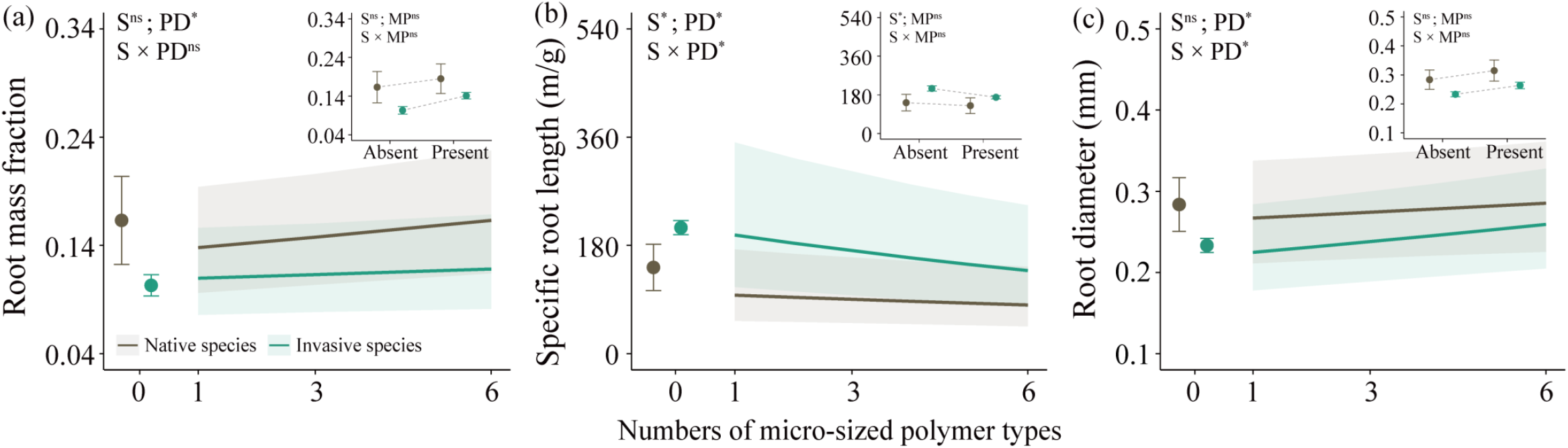
Root mass fraction (a), specific root length (b) and mean root diameter (c) of invasive and native species under the different numbers of micro-sized polymer types (0, 1, 3, 6) mixed into the substrate. In the main graph, points with error bars represent mean values ± SE in the absence of microplatic, and solid lines are the fitted relationships for the effect of polymer diversity, and the colored polygons delimit the 95% confidence intervals. Insets at the right upper corner of each graph show the effects of the absence *vs* presence of microplastic (averaged over 1, 3 and 6 polymer types). Significant parameters (i.e. ‘S’ [species invasion status], ‘PD’ [polymer diversity], ‘MP’ [microplastic pollution], ‘S × PD’ and ‘S × MP’) are indicted with asterisks (*), marginally significant ones are indicated with daggers (†), and non-significant ones are indicated with ‘ns’.

## Discussion

Our multi-species study found evidence that the presence of microplastic pollution suppressed growth of the invasive alien species stronger than for the overall smaller native species. However, the increased diversity of micro-sized polymer types resulted in stronger negative effects on plant growth. These negative effects of polymer diversity tended to be also stronger for invasive alien species than for native plants, possibly because with an increase in the number of polymer types, plants, and particularly invasive species, produced thicker roots, that are usually less absorptive.

We did not find a significant main effect of the presence of microplastic pollution —averaged over the three diversity levels— on plant growth. However, the significant interactive effect between invasion status of the species and microplastic pollution indicates that this does not apply to all species, and that the presence of microplastic pollution suppressed growth of invasive alien species more strongly than was the case for native species. This suggests that when the species would be grown in competition with each other, the presence of microplastic will change the dominance relations in the community. Lozano & Rillig (2020) grew artificial communities in the presence and absence of polyester fibers and indeed found changes in species dominance. The authors claimed that one of the species that increased in dominance, *Calamagrostis epigejos*, is invasive in Europe, and that microplastic pollution may contribute to its invasion success. However, although this species is expanding its distribution (Eichenberg et al. 2021) and dominance (Jandt et al. 2022) in Europe, it is native and thus not an invasive alien species to most of Eurasia (see its distribution in https://powo.science.kew.org). Our study, which explicitly compared multiple invasive and native plant species, suggests actually that microplastic pollution would decrease rather than increase invasion success of invasive species. In other words, compared to native species, invasive species took more advantage of soils that are not polluted, and thus that in the framework of Richards *et al*. (2006), they can be classified as ‘master of some’.

Our results showed that, in the presence of microplastic pollution, plants decreased their growth (i.e. total biomass) with increasing diversity of micro-sized polymer types. In other words, although soils polluted by different polymers may influence plant growth negatively or positively and to different extents (de Souza Machado *et al*. 2019), increased polymer diversity makes the effects of microplastic pollution on plants become more predictable. Our findings thus support the prediction of ‘sampling or synergistical effect’ hypothesis, and not the prediction of the ‘dilution effect’ hypothesis (Figure 1). Further analysis for the subset of plants grown in the absence or presence of each of the 12 micro-sized polymer types showed that only the polymers PBS, PCL and PLA significantly suppressed plant growth (Figure S1a). This is also supported by an additional experiment in which we tested the specific effect of each of the 12 micro-sized polymer types on growth of the native species *Eruca vesicaria* showed again that only the PBS, PCL and PLA strongly suppressed its growth (Figure S1b). With an increasing diversity of polymers, the probability increases that it contains at least one of these three polymers with strong negative effects. There could, however, also be synergistic effects of the different polymer types, but the observation that the 95% confidence interval (i.e. variance of effects) was much wider in the one-type-polymer treatments than in the six-types-polymer treatments (Figure 2) indicates that there likely was a strong sampling effect (Tilman, Lehman & Thomson 1997; Tilman & Lehman 2001).

In addition to the stronger negative effect of the presence of microplastic on invasive than on native species, invasive plants also tended to suffer more strongly from increasing polymer diversity than native species (Figure 2). This coincided with a positive effect of the number of polymer types on root thickness (i.e. a negative effect on root diameter and a positive effect on specific root length), which was also stronger for invasive than for native species (Figure 3b & c). Because thick roots are usually less absorptive than thin roots (Eghball & Maranville 1993; Ma *et al*. 2018), this could explain the reduced plant growth. Although plant could also improve their nutrient-uptake capacity by allocating more biomass to their root system (Bloom, Chapin & Mooney 1985; Liu & van Kleunen 2017), which they did in response to polymer diversity, we found no evidence that this response differed between invasive and native species. So, the stronger negative effect of polymer diversity on growth of invasive species might be due to the production of thick, less absorptive roots. In nature, the production of thicker roots could facilitate mycorrhization, and could then by outsourcing the nutrient uptake to the mycorrhizal fungi (Bergmann *et al*. 2020; Leifheit, Lehmann & Rillig 2021) increase instead of decrease growth. How mycorrhizal interactions of invasive and native plant species are affected by microplastic pollution and polymer diversity remains an open question.

Our study expands our understanding of the effects of multiple types of microplastic pollution on ecological processes. Thereby, it also supports the idea that the increasing number of simultaneously acting environmental changes could cause increasingly stronger directional changes in many ecological processes (Rillig *et al*. 2019b; Yang *et al*. 2022). For example, Zandalinas *et al*. (2011) showed that the accumulated impact of multifactorial stress combinations on plant growth and survival was detrimental. Therefore, even when we cannot study each possible multiway interaction among the many different global change factors or among the many types of polymers, our study and the few other studies mentioned above indicate that the number of simultaneously acting factors matter.

In conclusion, our multi-species experiment showed for the first time that the suppression of microplastic pollution of plant growth becomes stronger with increasing diversity of polymer types in the soil. Moreoever, we found tentative evidence that these negative effects of polymer diversity were greater for invasive species than for native plants, which could possibly be explained by changes in root thickness. While invasive plants in our study suffered more from the presence of micropastics and the diversity of polymers than natives did, invasive alien plants nevertheless produced still as much biomass as the natives did. Therefore, invasive alien plants might remain dominant, also in sites polluted with microplastics.

## Acknowledgements

We thank Xue Zhang, Zhengkuan Lu, Lu Xiao, Meng Hou, Lingxi Wang, Yimin Yan and Lichao Wang for assistance with the experiment. This work was financially supported by funding from the Chinese Academy of Sciences (Y9B7041001), the National Natural Science Foundation of China (32201341) and the Innovation Team Project of the Northeast Institute of Geography and Agroecology, Chinese Academy of Sciences (2022CXTD01).

## Author contributions

YL conceived the idea and designed the experiment. YF and KM performed the experiment. YF, MvK and YL analyzed the data. YF and YL wrote the draft of the manuscript, with major inputs from MvK.

## Data accessibility

Should the manuscript be accepted, the data supporting the results will be archived in Dryad and the data DOI will be included at the end of the article.

## Supporting information

**Table S1.**
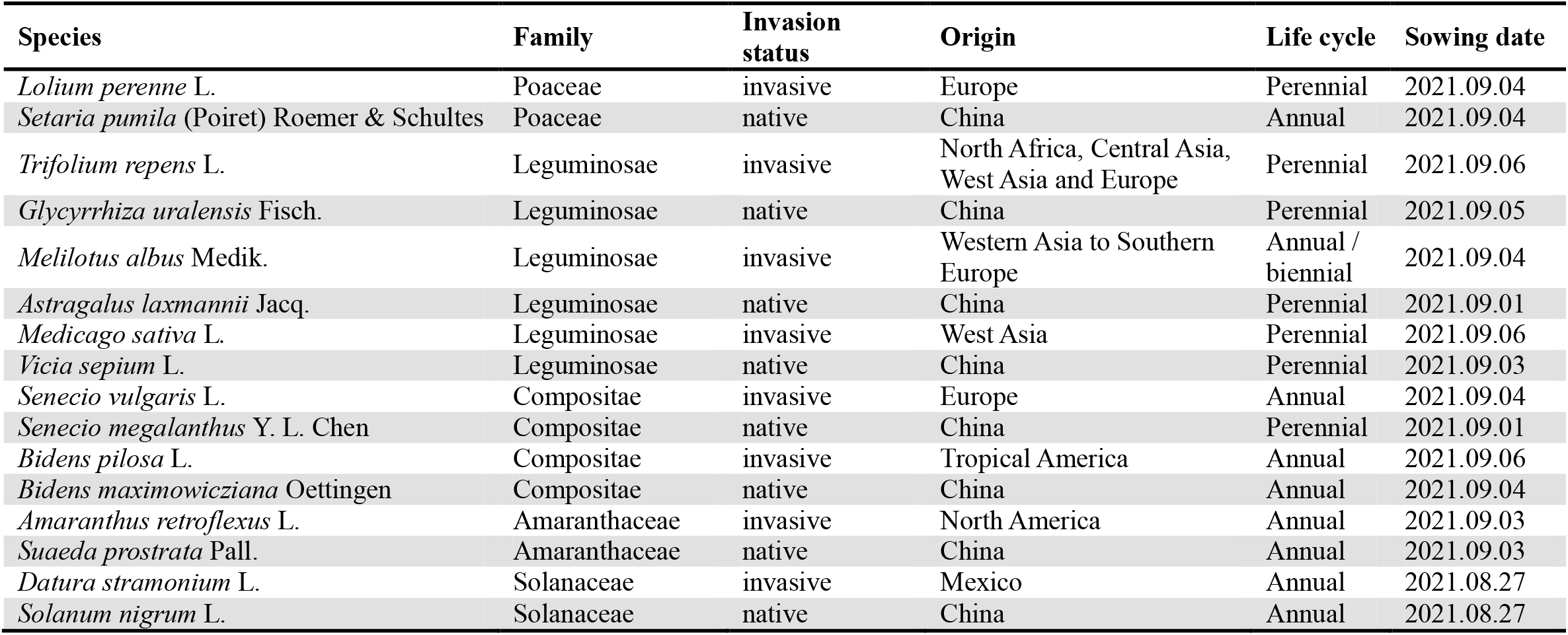
Details of the eight invasive and eight native species used in the experiment.

| Species | Family | Invasion status | Origin | Life cycle | Sowing date |
| --- | --- | --- | --- | --- | --- |
| *Lolium perenne* L. | Poaceae | invasive | Europe | Perennial | 2021.09.04 |
| *Setaria pumila* (Poiret) Roemer & Schultes | Poaceae | native | China | Annual | 2021.09.04 |
| *Trifolium repens* L. | Leguminosae | invasive | North Africa, Central Asia, West Asia and Europe | Perennial | 2021.09.06 |
| *Glycyrrhiza uralensis* Fisch. | Leguminosae | native | China | Perennial | 2021.09.05 |
| *Melilotus albus* Medik. | Leguminosae | invasive | Western Asia to Southern Europe | Annual / biennial | 2021.09.04 |
| *Astragalus laxmannii* Jacq. | Leguminosae | native | China | Perennial | 2021.09.01 |
| *Medicago sativa* L. | Leguminosae | invasive | West Asia | Perennial | 2021.09.06 |
| *Vicia sepium* L. | Leguminosae | native | China | Perennial | 2021.09.03 |
| *Senecio vulgaris* L. | Compositae | invasive | Europe | Annual | 2021.09.04 |
| *Senecio megalanthus* Y. L. Chen | Compositae | native | China | Perennial | 2021.09.01 |
| *Bidens pilosa* L. | Compositae | invasive | Tropical America | Annual | 2021.09.06 |
| *Bidens maximowicziana* Oettingen | Compositae | native | China | Annual | 2021.09.04 |
| *Amaranthus retroflexus* L. | Amaranthaceae | invasive | North America | Annual | 2021.09.03 |
| *Suaeda prostrata* Pall. | Amaranthaceae | native | China | Annual | 2021.09.03 |
| *Datura stramonium* L. | Solanaceae | invasive | Mexico | Annual | 2021.08.27 |
| *Solanum nigrum* L. | Solanaceae | native | China | Annual | 2021.08.27 |

**Table S2.**
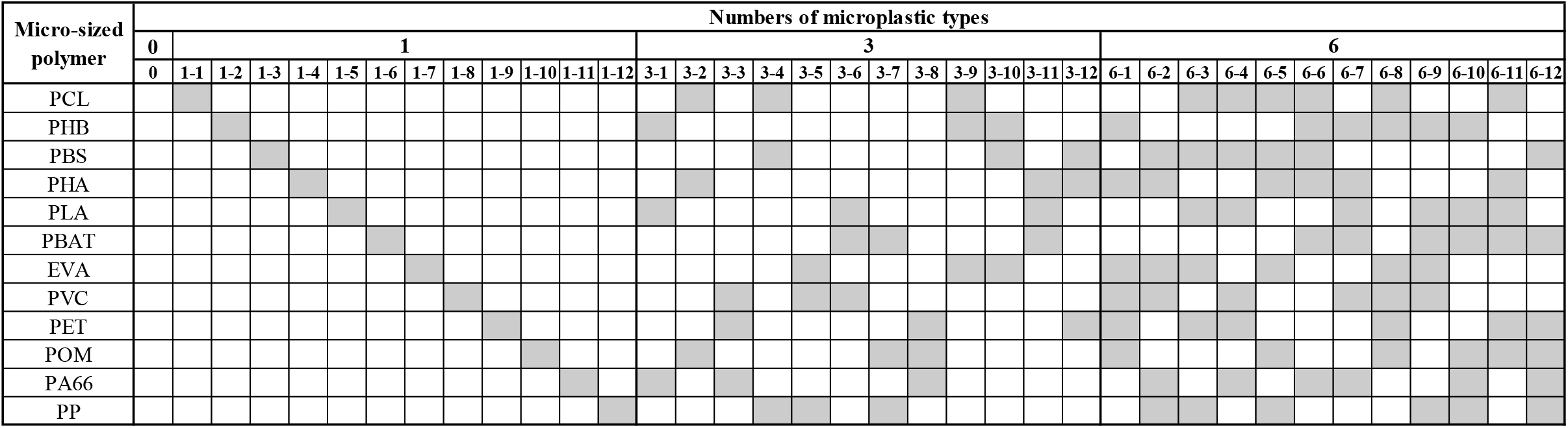
The different microplastic compositions used in the experiment. The number 0-6 represents the number of polymer types mixed into the substrate. The number from 1-1 to 6-12 represents each of the 36 different microplastic types or compositions used. Grey filling indicates that the polymer was present in the respective microplastic composition. See Figure 1c for the full names of the polymer types.

**Table S3.**
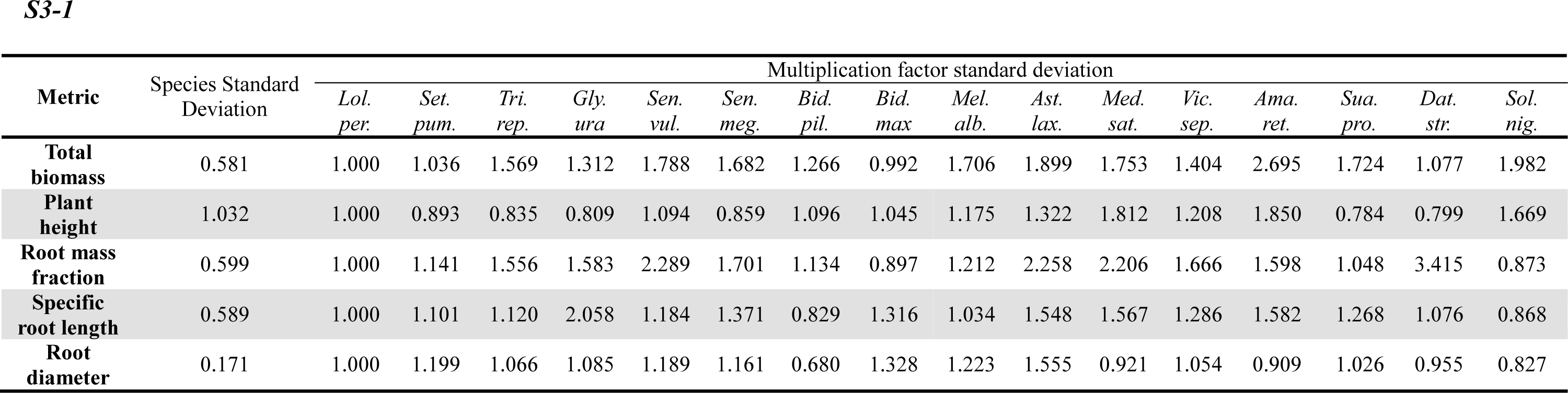
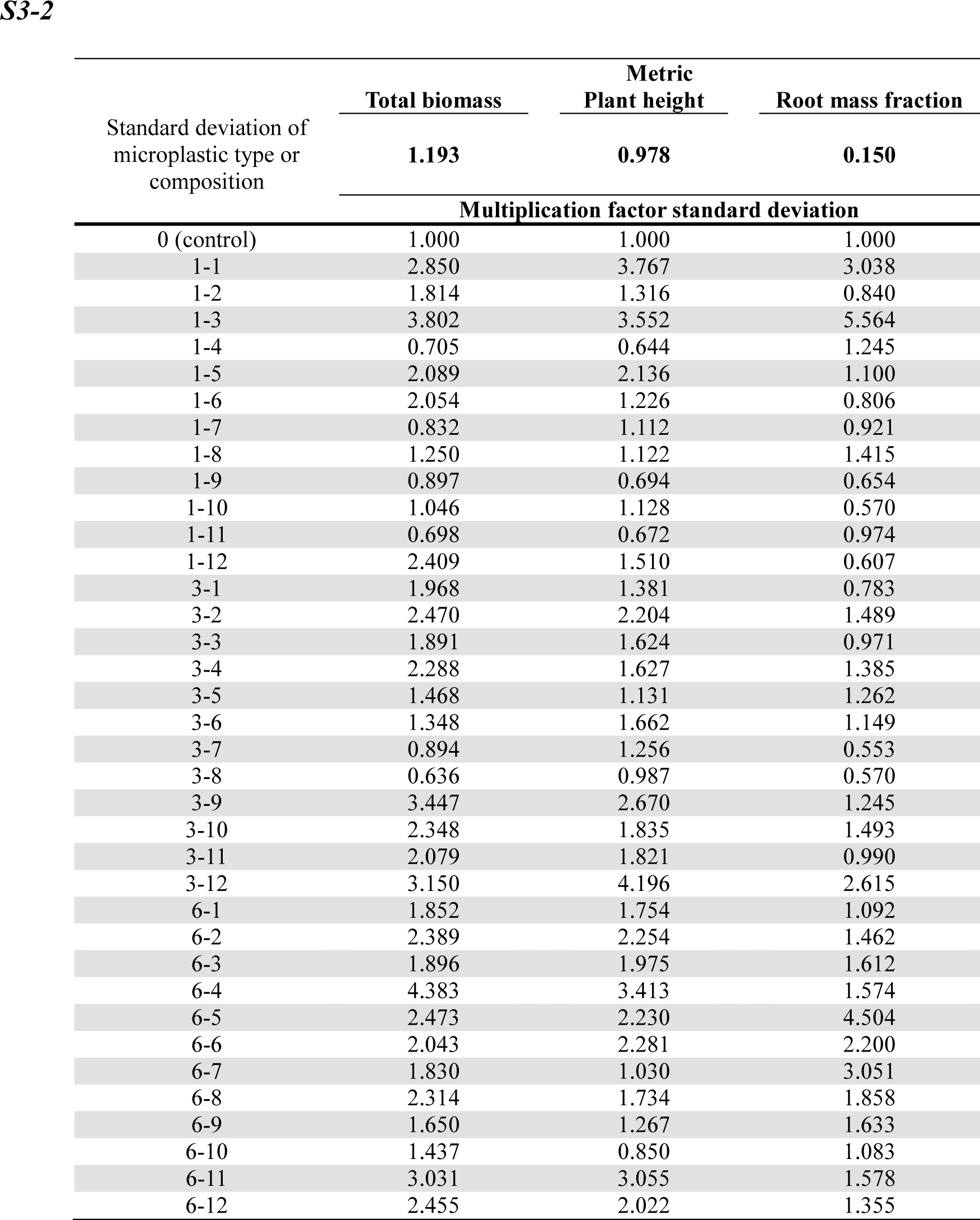
Standard deviations for individual species random effects for metrics analyzed with models with a Gaussian error distribution. The standard deviations given refer to the first species. For each species and microplastic composition, the given standard deviation should be multiplied by the multiplication factor. The names of the species in the table are abbreviated using the first three letters of the genus and the first three letters of the species epithet.

**Table S4.** Standard deviations for individual species random effects for metrics analyzed with models with a Gaussian error distribution. The standard deviations given refer to the first species. For each species and microplastic composition, the given standard deviation should be multiplied by the multiplication factor. The names of the species in the table are abbreviated using the first three letters of the genus and the first three letters of the species epithet.

**S4-1**
| **Metric** | Standard deviation of species | Multiplication factor standard deviation |  |  |  |  |  |  |  |  |  |  |  |  |  |  |  |
| --- | --- | --- | --- | --- | --- | --- | --- | --- | --- | --- | --- | --- | --- | --- | --- | --- | --- |
|  |  | *Lol. per.* | *Set. pum.* | *Tri. rep.* | *Gly. ura* | *Sen. vul.* | *Sen. meg.* | *Bid. pil.* | *Bid. max* | *Mel. alb.* | *Ast. lax.* | *Med. sat.* | *Vic. sep.* | *Ama. ret.* | *Sua. pro.* | *Dat. str.* | *Sol. nig.* |
| **Total biomass** | 0.616 | 1.000 | 0.839 | 1.231 | 1.086 | 2.141 | 1.490 | 1.341 | 0.996 | 1.749 | 1.659 | 1.637 | 1.439 | 2.683 | 1.546 | 1.244 | 2.152 |
| **Plant height** | 0.936 | 1.000 | 0.934 | 0.703 | 0.849 | 1.242 | 0.830 | 1.170 | 1.162 | 1.280 | 1.343 | 1.812 | 0.807 | 1.921 | 0.884 | 0.979 | 1.704 |
| **Root mass fraction** | 0.574 | 1.000 | 1.146 | 1.527 | 1.531 | 1.249 | 1.361 | 1.002 | 0.775 | 1.296 | 2.405 | 2.380 | 1.748 | 1.418 | 1.146 | 1.337 | 0.990 |
| **Specific root length** | 0.602 | 1.000 | 1.002 | 1.038 | 1.840 | 1.205 | 1.183 | 0.758 | 1.300 | 1.031 | 1.426 | 1.355 | 1.165 | 1.482 | 1.249 | 1.038 | 0.730 |
| **Root diameter** | 0.172 | 1.000 | 1.067 | 1.021 | 0.986 | 1.120 | 1.087 | 0.605 | 1.179 | 1.213 | 1.287 | 0.868 | 0.912 | 0.678 | 1.018 | 0.766 | 0.740 |

| Standard deviation of<br>microplastic type or<br>composition | Total biomass | Metric<br>Plant height | Root mass fraction |
| --- | --- | --- | --- |
|  | 1.102 | 0.907 | 0.113 |
|  | Multiplication factor standard deviation |  |  |
| 1-1 | 1.000 | 1.000 | 1.000 |
| 1-2 | 0.464 | 0.360 | 0.146 |
| 1-3 | 1.072 | 0.905 | 0.818 |
| 1-4 | 0.243 | 0.215 | 0.189 |
| 1-5 | 0.603 | 0.549 | 0.160 |
| 1-6 | 0.619 | 0.381 | 0.162 |
| 1-7 | 0.286 | 0.361 | 0.144 |
| 1-8 | 0.460 | 0.365 | 0.214 |
| 1-9 | 0.312 | 0.191 | 0.125 |
| 1-10 | 0.352 | 0.330 | 0.112 |
| 1-11 | 0.235 | 0.194 | 0.173 |
| 1-12 | 0.647 | 0.481 | 0.105 |
| 3-1 | 0.615 | 0.398 | 0.121 |
| 3-2 | 0.762 | 0.583 | 0.257 |
| 3-3 | 0.609 | 0.453 | 0.152 |
| 3-4 | 0.697 | 0.412 | 0.220 |
| 3-5 | 0.408 | 0.309 | 0.199 |
| 3-6 | 0.401 | 0.542 | 0.168 |
| 3-7 | 0.346 | 0.405 | 0.078 |
| 3-8 | 0.231 | 0.272 | 0.087 |
| 3-9 | 1.034 | 0.737 | 0.184 |
| 3-10 | 0.626 | 0.472 | 0.221 |
| 3-11 | 0.567 | 0.516 | 0.166 |
| 3-12 | 0.920 | 1.109 | 0.408 |
| 6-1 | 0.612 | 0.467 | 0.204 |
| 6-2 | 0.661 | 0.625 | 0.216 |
| 6-3 | 0.565 | 0.558 | 0.250 |
| 6-4 | 1.196 | 0.888 | 0.284 |
| 6-5 | 0.708 | 0.606 | 0.759 |
| 6-6 | 0.570 | 0.620 | 0.340 |
| 6-7 | 0.552 | 0.277 | 0.440 |
| 6-8 | 0.696 | 0.481 | 0.280 |
| 6-9 | 0.514 | 0.328 | 0.258 |
| 6-10 | 0.416 | 0.235 | 0.205 |
| 6-11 | 0.882 | 0.763 | 0.230 |
| 6-12 | 0.661 | 0.516 | 0.225 |

**Figure S1.**
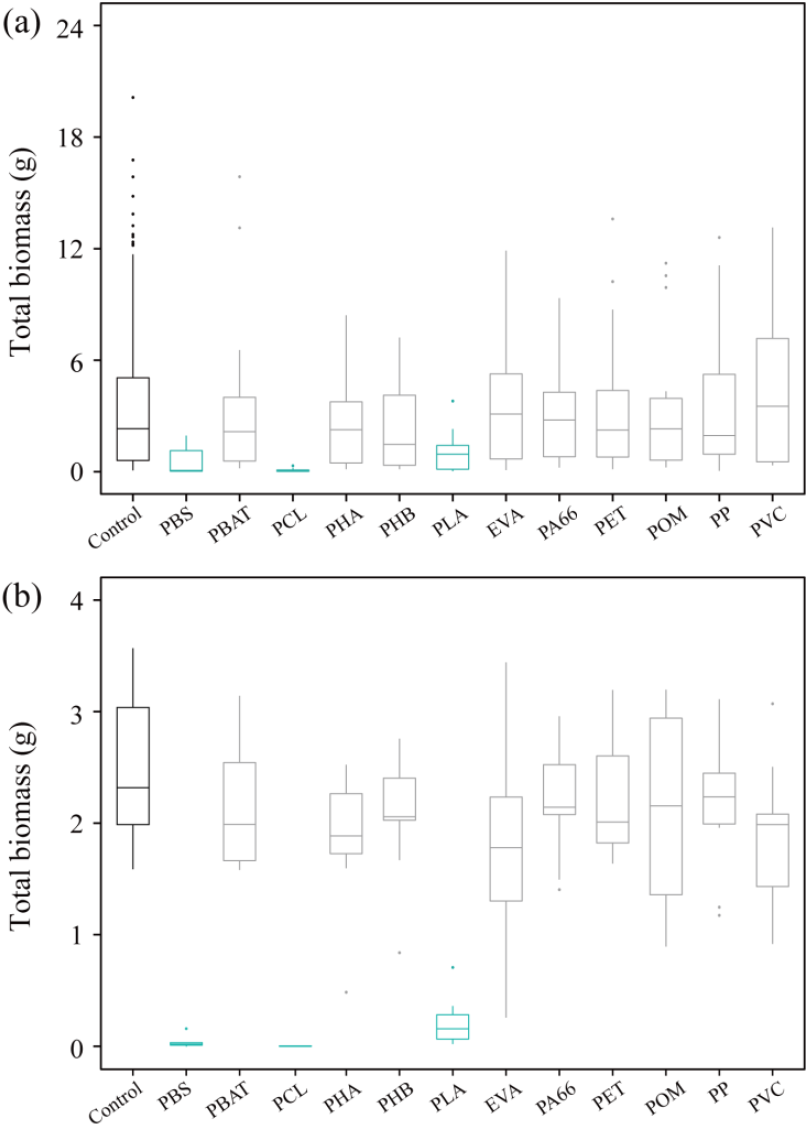
Box plots showing total biomass of plants (averaged across 16 species) in present study (a) and species *Eruca vesicaria* (averaged across ten replicates) grown in the additionally experiment (b). The horizontal line inside the box is the median value, and the upper and lower bounds of the box indicate the interquartile range (i.e. the 75^th^ and 25^th^ percentiles, respectively). Whiskers represent data within 1.5 times the interquartile range, and individual dots are outliers. The grey boxes indicate no significant differences (*p* > 0.05) between the specific microplastic type and the control treatment (i.e. without microplatic), while the green boxes indicate significant differences (*p* < 0.05) between the specific microplastic type and the control treatment. The significance tests were done by linear regression models using the lm function in the R package ‘stats’ and summary function in the R package ‘base’. See Figure 1c for the full names of the polymer types.

## Reference

Baho, D.L., Bundschuh, M. & Futter, M.N. (2021) Microplastics in terrestrial ecosystems: Moving beyond the state of the art to minimize the risk of ecological surprise. Global Change Biology, 27, 3969–3986.

Barnes, D.K., Galgani, F., Thompson, R.C. & Barlaz, M. (2009) Accumulation and fragmentation of plastic debris in global environments. Philosophical transactions of the royal society B: biological sciences, 364, 1985–1998.

Bergmann, J., Weigelt, A., van der Plas, F., Laughlin, D.C., Kuyper, T.W., Guerrero-Ramirez, N., Valverde-Barrantes, O.J., Bruelheide, H., Freschet, G.T., Iversen, C.M., Kattge, J., McCormack, M.L., Meier, I.C., Rillig, M.C., Roumet, C., Semchenko, M., Sweeney, C.J., van Ruijven, J., York, L.M. & Mommer, L. (2020). The fungal collaboration gradient dominates the root economics space in plants. Science Advances, 6, eaba3756.

Bloom, A.J., Chapin, F.S. & Mooney, H.A. (1985) Resource limitation in plants—an economic analogy. Annual review of ecology and systematics, 363–392.

Boots, B., Russell, C.W. & Green, D.S. (2019) Effects of microplastics in soil ecosystems: above and below ground. Environmental Science & Technology, 53, 11496–11506.

Bosker, T., Bouwman, L.J., Brun, N.R., Behrens, P. & Vijver, M.G. (2019) Microplastics accumulate on pores in seed capsule and delay germination and root growth of the terrestrial vascular plant Lepidium sativum. Chemosphere, 226, 774–781.

Chytrý M, Jarosik V, Pysek P, Hájek O, Knollová I, Tichý L, Danihelka J. (2008) Separating habitat invasibility by alien plants from the actual level of invasion. Ecology 89, 1541–1553.

de Souza Machado, A.A., Kloas, W., Zarfl, C., Hempel, S. & Rillig, M.C. (2018) Microplastics as an emerging threat to terrestrial ecosystems. Global Change Biology, 24, 1405–1416.

de Souza Machado, A.A., Lau, C.W., Kloas, W., Bergmann, J., Bachelier, J.B., Faltin, E., Becker, R., Görlich, A.S. & Rillig, M.C. (2019) Microplastics can change soil properties and affect plant performance. Environmental Science & Technology, 53, 6044–6052.

Eghball, B. & Maranville, J.W. (1993) Root development and nitrogen influx of corn genotypes grown under combined drought and nitrogen stresses. Agronomy Journal, 85, 147–152.

Eichenberg, D., Bowler, D.E., Bonn, A., Bruelheide, H., Grescho, V., Harter, D., Jandt, U., May, R., Winter, M. & Jansen, F. (2021), Widespread decline in Central European plant diversity across six decades. Global Change Biology, 27, 1097–1110.

Fox, J. & Weisberg, S. (2019) An R Companion to Applied Regression, Third edition. Sage, Thousand Oaks CA.

Fuller, S. & Gautam, A. (2016) A procedure for measuring microplastics using pressurized fluid extraction. Environmental Science & Technology, 50, 5774–5780.

Geyer, R., Jambeck, J.R. & Law, K.L. (2017) Production, use, and fate of all plastics ever made. Science advances, 3, e1700782.

Jambeck, J.R., Geyer, R., Wilcox, C., Siegler, T.R., Perryman, M., Andrady, A., Narayan, R. & Law, K.L. (2015) Plastic waste inputs from land into the ocean. Science, 347, 768–771.

Jandt, U., Bruelheide, H., Jansen, F., Bonn, A., Grescho, V., Klenke, R.A., Sabatini, F.M., Bernhardt-Römermann, M., Blüml, V., Dengler, J., Diekmann, M., Doerfler, I., Döring, U., Dullinger, S., Haider, S., Heinken, T., Horchler, P., Kuhn, G., Lindner, M., Metze, K.. Müller, N., Naaf, T., Peppler-Lisbach, C., Poschlod, P., Roscher, C., Rosenthal, G., Rumpf, S.B., Schmidt, W., Schrautzer, J., Schwabe, A., Schwartze, P., Sperle, T., Stanik, N., Storm, C., Voigt, W., Wegener, U., Wesche, K., Wittig, B. & Wulf, M. (2022). More losses than gains during one century of plant biodiversity change in Germany. Nature, 611, 512–518

Leifheit, E.F., Lehmann, A. & Rillig, M.C. (2021) Potential effects of microplastic on arbuscular mycorrhizal fungi. Frontiers in Plant Science, 12, 626709.

Lewis, S.L. & Maslin, M.A. (2015) Defining the anthropocene. Nature, 519, 171–180.

Liu, Y. & van Kleunen, M. (2017) Responses of common and rare aliens and natives to nutrient availability and fluctuations. Journal of Ecology, 105, 1111–1122.

Loreau, M. & Hector, A. (2001). Partitioning selection and complementarity in biodiversity experiments. Nature, 412, 72–76.

Lozano, Y.M., Aguilar-Trigueros, C.A., Onandia, G., Maaß, S., Zhao, T. & Rillig, M.C. (2021a) Effects of microplastics and drought on soil ecosystem functions and multifunctionality. Journal of Applied Ecology, 58, 988–996.

Lozano, Y.M., Lehnert, T., Linck, L.T., Lehmann, A. & Rillig, M.C. (2021b) Microplastic shape, polymer type, and concentration affect soil properties and plant biomass. Frontiers in Plant Science, 12, 616645.

Lozano, Y.M. & Rillig, M.C. (2020) Effects of microplastic fibers and drought on plant communities. Environmental Science & Technology 54, 6166–6173.

Luo, Y., Li, L., Feng, Y., Li, R., Yang, J., Peijnenburg, W.J. & Tu, C. (2022) Quantitative tracing of uptake and transport of submicrometre plastics in crop plants using lanthanide chelates as a dual-functional tracer. Nature nanotechnology, 17, 424–431.

Ma, J.S. (2020). tAlien Invasive Flora of China. Shanghai Jiao Tong University Press.

Ma, Z., Guo, D., Xu, X., Lu, M., Bardgett, R.D., Eissenstat, D.M., McCormack, M.L. & Hedin, L.O. (2018) Evolutionary history resolves global organization of root functional traits. Nature, 555, 94–97.

Meng, F., Yang, X., Riksen, M., Xu, M. & Geissen, V. (2021) Response of common bean (Phaseolus vulgaris L.) growth to soil contaminated with microplastics. Science of the Total Environment, 755, 142516.

Pinheiro, J., Bates, D. & R Core Team (2020) nlme: linear and nonlinear mixed effects models. R package version 3.1-148.

R Core Team (2020) R: A language and environment for statistical computing. R Foundation for Statistical Computing, Vienna, Austria. URL https://www.R-project.org/.

Richards, C.L., Bossdorf, O., Muth, N.Z., Gurevitch, J. & Pigliucci, M. (2006) Jack of all trades, master of some? On the role of phenotypic plasticity in plant invasions. Ecology Letters, 9, 981–993.

Rillig, M.C. (2018) Microplastic disguising as soil carbon storage. Environmental science & technology, 52, 6079–6080.

Rillig, M.C. & Lehmann, A. (2020) Microplastic in terrestrial ecosystems. Science, 368, 1430–1431.

Rillig, M.C., Lehmann, A., de Souza Machado, A.A. & Yang, G. (2019a) Microplastic effects on plants. New Phytologist, 223, 1066–1070.

Rillig, M.C., Ryo, M., Lehmann, A., Aguilar-Trigueros, C.A., Buchert, S., Wulf, A., Iwasaki, A., Roy, J. & Yang, G. (2019b) The role of multiple global change factors in driving soil functions and microbial biodiversity. Science, 366, 886–890.

Seebens, H., Blackburn, T.M., Dyer, E.E., Genovesi, P., Hulme, P.E., Jeschke, J.M., Pagad, S., Pyšek, P., van Kleunen, M., Winter, M., Ansong, M., Arianoutsou, M., Bacher, S., Blasius, B., Brockerhoff, E.G., Brundu, G., Capinha, C., Causton, C.E., Celesti-Grapow, L., Dawson, W., Dullinger, S., Economo, E.P., Fuentes, N., Guénard, B., Jäger, H., Kartesz, J., Kenis, M., Kühn, I., Lenzner, B., Liebhold, A.M., Mosena, A., Moser, D., Nentwig, W., Nishino, M., Pearman, D., Pergl, J., Rabitsch, W., Rojas-Sandoval, J., Roques, A., Rorke, S., Rossinelli, S., Roy, H.E., Scalera, R., Schindler, S., Štajerová, K., Tokarska-Guzik, B., Walker, K., Ward, D.F., Yamanaka, T. & Essl, F. (2018) Global rise in emerging alien species results from increased accessibility of new source pools. Proceedings of the National Academy of Sciences, 115, E2264–E2273.

Sun, X.D., Yuan, X.Z., Jia, Y., Feng, L.J., Zhu, F.P., Dong, S.S., Liu, J., Kong, X., Tian, H., Duan, J.L., Ding, Z., Wang, S.G. & Xing, B. (2020) Differentially charged nanoplastics demonstrate distinct accumulation in Arabidopsis thaliana. Nature nanotechnology, 15, 755–760.

Thompson, R.C., Olsen, Y., Mitchell, R.P., Davis, A., Rowland, S.J., John, A.W., McGonigle, D. & Russell, A.E. (2004) Lost at sea: where is all the plastic? Science, 304, 838.

Tilman, D., Lehman, C.L. & Thomson, K.T. (1997) Plant diversity and ecosystem productivity: Theoretical considerations. Proceedings of the National Academy of Sciences, 94, 1857–1861.

Tilman, D. & Lehman, C.L. (2001) Biodiversity, composition and ecosystem processes: theory and concepts. In: Kinzig, A.P., Pacala, S.W. & Tilman, D. (eds.) The Functiona Consequences of Biodiversity: Empirical Progress and Theoretical Extensions. Monographs in Population Biology 33, pp. 9–41, Princeton University Press, Princeton and Oxford.

van Kleunen, M., Brumer, A., Gutbrod, L. & Zhang, Z. (2020) A microplastic used as infill material in artificial sport turfs reduces plant growth. Plants, People, Planet, 2, 157–166.

van Kleunen, M., Dawson, W., Essl, F., Pergl, J., Winter, M., Weber, E., Kreft, H., Weigelt, P., Kartesz, J., Nishino, M., Antonova, L.A., Barcelona, J.F., Cabezas, F.J., Cárdenas, D., Cárdenas-Toro, J., Castaño, N., Chacón, E., Chatelain, C., Ebel, A.L., Figueiredo, E., Fuentes, N., Groom, Q.J., Henderson, L., Inderjit Kupriyanov, A., Masciadri, S., Meerman, J., Morozova, O., Moser, D., Nickrent, D.L., Patzelt, A., Pelser, P.B., Baptiste, M.P., Poopath, M., Schulze, M., Seebens, H., Shu, W.S., Thomas, J., Velayos, M., Wieringa, J.J. & Pyšek, P. (2015) Global exchange and accumulation of non-native plants. Nature, 525, 100–103.

Wei, X.F., Nilsson, F., Yin, H. & Hedenqvist, M.S. (2021) Microplastics originating from polymer blends: an emerging threat? Environmental Science & Technology, 55, 4190–4193.

Xu, C., Zhang, B., Gu, C., Shen, C., Yin, S., Aamir, M. & Li, F. (2020) Are we underestimating the sources of microplastic pollution in terrestrial environment? Journal of Hazardous Materials, 400, 123228.

Yang, G., Ryo, M., Roy, J., Lammel, D.R., Ballhausen, M.B., Jing, X., Zhu, X. & Rillig, M.C. (2022) Multiple anthropogenic pressures eliminate the effects of soil microbial diversity on ecosystem functions in experimental microcosms. Nature communications, 13, 4260.

Yang, W., Cheng, P., Adams, C.A., Zhang, S., Sun, Y., Yu, H. & Wang, F. (2021) Effects of microplastics on plant growth and arbuscular mycorrhizal fungal communities in a soil spiked with ZnO nanoparticles. Soil Biology and Biochemistry, 155, 108179.

Zandalinas, S.I., Sengupta, S., Fritschi, F.B., Azad, R.K., Nechushtai, R. & Mittler, R. (2021) The impact of multifactorial stress combination on plant growth and survival. New Phytologist, 230, 1034–1048.

Zuur, A.F., Ieno, E.N., Walker, N.J., Saveliev, A.A. & Smith, G.M. (2009) Mixed effects models and extensions in ecology with R. Springer Science & Business Media.

